# Analysis of p67 allelic sequences reveals a subtype of allele type 1 unique to buffalo-derived *Theileria parva* parasites from southern Africa

**DOI:** 10.1101/2020.03.25.007583

**Authors:** Lubembe D. Mukolwe, David O. Odongo, Charles Byaruhanga, Louwtjie P. Snyman, Kgomotso P. Sibeko-Matjila

## Abstract

East Coast fever (ECF) and Corridor disease (CD) caused by cattle- and buffalo-derived *T. parva* respectively are the most economically important tick-borne diseases of cattle in the affected African countries. The p67 gene has been evaluated as a recombinant subunit vaccine against East Coast fever (ECF), and for discrimination of *T. parva* parasites causing ECF and Corridor disease (CD). The p67 allele type 1 was first identified in cattle-derived *T. parva* parasites from east Africa, where parasites possessing this allele type have been associated with ECF. Subsequent characterization of buffalo-derived *T. parva* parasites from South Africa where ECF was eradicated, revealed the presence of a similar allele type, raising concerns as to whether or not allele type 1 from parasites from the two regions is identical. A 900 bp central fragment of the gene encoding p67 was PCR amplified from *T. parva* DNA extracted from blood collected from cattle and buffalo in South Africa, Mozambique, Kenya, Tanzania and Uganda, followed by DNA sequence analysis. Four p67 allele types previously described were identified. A subtype of p67 allele type 1 was identified in parasites from clinical cases of CD and buffalo from southern Africa. Notably, p67 allele type 1 sequences from parasites associated with ECF in East Africa and CD in Kenya were identical. Analysis of two p67 B-cell epitopes (TpM12 and AR22.7) revealed amino acid substitutions in allele type 1 from buffalo-derived *T. parva* parasites from southern Africa. However, both epitopes were conserved in allele type 1 from cattle- and buffalo-derived *T. parva* parasites from East Africa. These findings reveal detection of a subtype of p67 allele type 1 associated with *T. parva* parasites transmissible from buffalo to cattle in southern Africa.

## Introduction

Theileriosis is a widespread tick-transmitted protozoal disease of wildlife and domestic animals caused by an apicomplexan parasite of the genus *Theileria* (reviewed in 1). In eastern, central and southern Africa, cattle theileriosis is commonly caused by *Theileria parva* which occurs naturally in the African buffalo (*Syncerus caffer*) that is an asymptomatic carrier (2). *Theileria parva* causes fatal classical East Coast fever (ECF) (3, 4), Corridor disease (5, 6) and January disease (7) occurring in different African countries. This parasite is mainly transmitted by a three-host brown ear tick *Rhipicephalus appendiculatus*, although *Rhipicephalus zambeziensis* and *Rhipicephalus duttoni* are also possible vectors (3, 8, 9).

East Coast fever is estimated to result in an economic loss of about USD 300 due to death of approximately one million cattle annually in the affected countries (reviewed in 1). A live trivalent sporozoite vaccine for control of ECF was developed (10) and has been successfully adopted for use in East Africa (11, 12). Although this vaccine does not confer protection against buffalo-derived *T. parva* in Kenya (13), studies in northern Tanzania have suggested that the vaccine is effective in areas where cattle co-graze with buffalo (14, 15). Efforts to develop a recombinant vaccine based on the antigen genes, to confer protection against cattle- and buffalo-derived *T. parva*, have been pursued for the last four decades (16), however, up to date, this has not been successful. Immunity to *T. parva* infections is mainly cell-mediated involving CD8+ cytotoxic T lymphocytes which recognize parasite peptides encoded by schizont genes and presented by MHC Class I molecules (17, 18). In addition, it has been demonstrated that antigens recognized by monoclonal antibodies may induce an antibody-mediated immune protection (19).

*Theileria parva* schizont and sporozoite antigen genes encoding proteins recognized by the host’s immune system have been identified (20–22) and characterized (19, 23–28). The gene encoding the sporozoite antigen, p67, has been explored for the development of a recombinant vaccine. Vaccination of cattle using a subunit vaccine based on the recombinant versions of p67 demonstrated a reduction in disease incidence by approximately 70% in the laboratory (29, 30) and about 30% under field tick challenge (31). Recent attempts have been made to improve the vaccination regimen by modifying the antigen preparation, dosage or adjuvant systems (32).

Previous studies on the sequence diversity of the central variable region of the p67 gene in cattle- and buffalo-derived *T. parva* parasites from Kenya (21, 33, 34) and South Africa (26) revealed four groups of related p67 alleles referred to as types 1, 2, 3 and 4. The different allele types are distinguishable based on two indels; a 129 bp insert absent in allele type 1 and present in type 2 (21), and a 174 bp insert absent in allele type 3 and present in allele type 4 (26). Five distinct p67 B-cell epitopes recognized by murine monoclonal antibodies have been identified (19) with sequence polymorphism occurring in buffalo-derived *T. parva* parasites (19, 33).

Cattle-derived *T. parva* parasites are known to have identical p67 allele type 1 sequences, and parasites possessing this allele type have been associated with classical ECF (21). Recently, it was established that buffalo-derived *T. parva* parasites implicated in Corridor disease in Kenya also possess p67 allele type 1 (34) that is identical to that from cattle-derived *T. parva* parasites. In South Africa, p67 allele type 1 similar to that identified in East Africa was identified from buffalo-derived *T. parva* parasites (26). Collectively, these studies have demonstrated that parasites possessing p67 allele type 1 are responsible for ECF in East Africa (21), and are also involved in Corridor disease in Kenya (34) and South Africa (26). However, in South Africa, there are no reports of ECF since its eradication in the 1950s, yet p67 allele type 1 has been detected in buffalo-derived *T. parva* parasites (26). One possible explanation to this phenomenon is that parasites associated with *T. parva* infections in cattle in East and southern Africa differ based on p67 allele type 1 sequence. To investigate this, we performed a comparative sequence analysis of the p67 gene in cattle- and buffalo-derived *T. parva* field parasites from East and southern Africa.

## Materials and Methods

### Ethics Statement

This study was approved by the Animal Ethics Committee of the Faculty of Veterinary Science, University of Pretoria (AEC Certificate # V080-16). Additional approvals including Section 20 and import permits were obtained from the Department of Agriculture, Forestry and Fisheries, South Africa.

### Sample collection and detection of *T. parva*

Blood samples were collected from cattle and buffalo from South Africa, Mozambique, Kenya, Tanzania and Uganda, in both present and previous studies, as indicated in Table 1. In addition, two DNA samples were obtained from Katete and Chitongo *T. parva* isolates from Zambia. DNA was extracted from the blood samples using the DNeasy® Blood and Tissue kit (Qiagen, Hilden, Germany), according to the manufacturer’s protocol. However, elution was done in 100 μl instead of the recommended 200 μl to increase the concentration of extracted DNA. Extracted DNA was stored at 4°C and – 20°C for short- and long-term storage respectively until further analysis. Detection of *T. parva* genomic DNA was done using a *T. parva*-specific hybridization probe-based real-time PCR assay targeting the 18S rRNA gene (35).

**Table 1.**
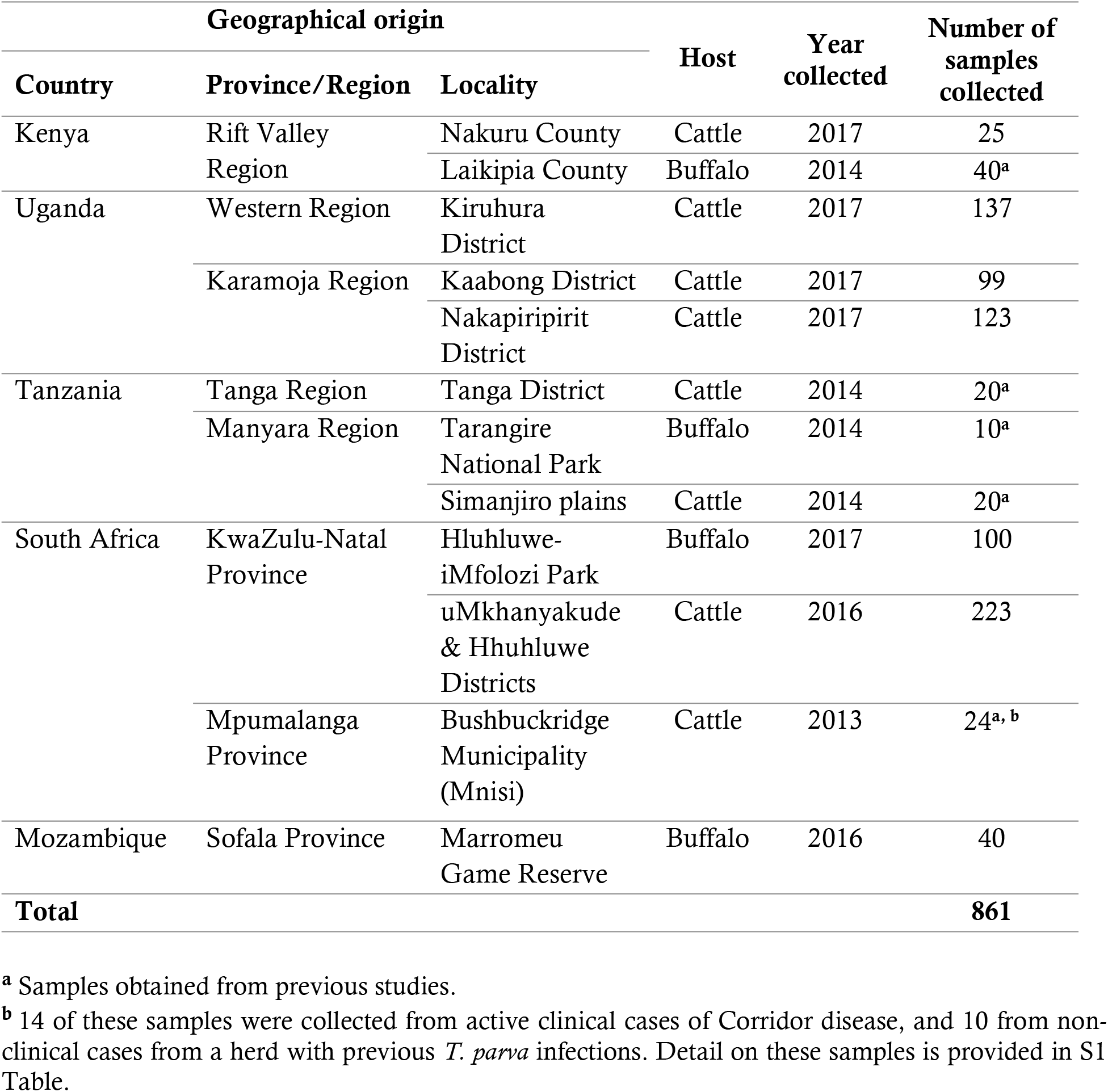
The geographical origin of blood samples collected from East and southern Africa

### PCR amplification of the gene encoding the p67 antigen

PCR amplification targeting the 900 bp variable region of the p67 encoding gene was performed on 232 *T. parva* positive DNA samples, using the primer pair IL613 (5’-ACAAACACAATCCCAAGTTC-3’) and IL792 (5’-CCTTTACTACGTTGGCG-3’) (21) and the 2X Phusion™ Flash High-Fidelity PCR Master Mix (ThermoFisher Scientific™, Waltham MA, USA) containing Phusion Flash II DNA polymerase which has proof-reading activity (36). At least 50 ng of DNA and 10 pmol of each primer were used in a total reaction volume of 12.5 μl. The amplification conditions were as previously described by Nene, Musoke (21), with some modifications in accordance with the Phusion Flash High-Fidelity PCR Master Mix conditions. Thus, the initial denaturation was done at 98°C for 10 seconds (s), followed by 30 cycles of denaturation at 98°C for 1 s, annealing at 57°C for 5 s and extension at 72°C for 10 s, then one cycle for the final extension step at 72°C for 1 minute (min). Samples that failed to amplify in the primary reaction were re-amplified in the second PCR using 0.5 μl of the primary PCR product as DNA template, and the same amplification conditions except that the amplification cycles were reduced to 20. PCR products were resolved by gel electrophoresis using 2% agarose stained with ethidium bromide.

### Cloning and Sanger sequencing

The PCR products were purified using the QIAquick^®^ PCR Purification Kit (Qiagen, Hilden, Germany) according to the manufacturer’s protocol, except that the final elution was done in 25 μl instead of the recommended 50 μl to increase the concentration of DNA. Purified PCR products were ligated into pJET1.2/blunt cloning vector using the ClonJET™ PCR cloning Kit (ThermoFisher Scientific™, Waltham, MA USA), followed by transformation of JM109 *E. coli* competent cells (Zymo Research, Tustin, USA). Recombinant clones were confirmed by colony PCR performed in a 20 μl reaction volume consisting of 2X DreamTaq Green PCR Master Mix (ThermoFisher Scientific™, Waltham, MA USA), forward and reverse pJET primers each at 4 pmol. The cycling conditions were as follows; an initial denaturation at 95°C for 3 min, followed by 25 cycles of denaturation at 94°C for 30 s, annealing at 60°C for 30 s and extension at 72°C for 1 min, then one cycle for the final extension step at 72°C for 1 min. The PCR products were analyzed by gel electrophoresis using 2% agarose in 1X TAE running buffer. Colony PCR products were purified using the QIAquick^®^ PCR Purification Kit (Qiagen, Hilden, Germany) following manufacturer’s protocol. Bidirectional sequencing was performed using pJET primers on ABI 3500XL Genetic Analyzer, POP7™ (ThermoFisher Scientific™, Waltham MA, USA) at INQABA Biotechnologies, South Africa.

### Sequence analysis

Raw p67 sequences were confirmed using the Basic Local Alignment Search Tool (BLAST), and sequence assembly and editing were done using the CLC Main Workbench version 8.0 (Qiagen, Hilden, Germany). Multiple sequence alignment of consensus sequences, together with reference sequences (Supp. Table S6), was done using the online version 7 of MAFFT (37) applying the default parameters (http://mafft.cbrc.jp/alignment/server/). Estimation of the effect of amino acid substitutions within the epitope regions was done using SIFT predictions (38) where a probability score >0.05 was predicted to be tolerant.

### Phylogenetic analysis

The aligned sequence matrix was truncated, manually viewed and edited using MEGA version 7 (39), where the final matrix was used for all subsequent analyses. Format changes on the sequence matrix for different analyses was achieved using the HIV sequence database online format converter tool (http://www.hiv.lanl.gov/). A data-display network (neighbor-joining network) was generated using SplitsTree version 4 (40), where all characters and uncorrected p-distances were used, and bootstrap support calculated from 1000 replicates. Evidence of recombination was determined by performing a phi-test in SplitsTree. The jModelTest version 2 (41) executed on the Cipres Science Gateway (https://www.phylo.org/portal2/) was used for model estimation, where Akaike Information Criterion and Bayesian Information Criterion were evaluated. Maximum Likelihood (ML) analysis was performed using RAxML version 8 (42) employing the GTR model, where gamma distribution and invariable sites were included as prescribed by the model estimation. No partitioning or outgroups were specified, and analysis was initiated using a random starting tree. The autoMRE bootstrapping function in RAxML was invoked for calculating bootstrap support. Bayesian inference was performed using MrBayes version 3 (43), applying the same model specifications as in the ML analysis. Two Markov Chain Monte Carlo (MCMC) chains were searched for 10 million iterations, saving every 1000th tree. After collating the saved trees, the first 15% were discarded as burn-in and posterior probabilities were calculated from the remaining trees. Parameter stabilization and the effective sample size (ESS) value were assessed using Tracer version 1.6 (44). The trees were viewed in FigTree version 1.4.3 (http://tree.bio.ed.ac.uk/software/figtree/) and exported for graphical modification in Corel Paintshop Pro X8. Bootstrap support greater than 70, and posterior probabilities greater than 0.9 were displayed as branch support on the ML topology.

## Results

### Detection of *T. parva* genomic DNA

Out of the 861 DNA samples (buffalo = 190, cattle = 671) extracted from whole blood, 306 consisting of 166 and 140 samples from cattle and buffalo, respectively, were detected as positive for *T. parva*.

### Size discrimination of p67 PCR amplicons

PCR amplification of the p67 gene targeting the central variable region was successfully done on 128 of the 232 *T. parva* positive DNA samples selected for analysis (Table 2). Amplicons of four varying fragment sizes were detected as single or multiple bands of 800 900 bp, 1000 bp and 1100 bp previously associated with allele types 3, 1, 4 and 2, respectively (21, 26). Most of the *T. parva* positive DNA samples from buffalo as well as clinical cases of Corridor disease from South Africa generated multiple amplicon profiles representing all four p67 fragment sizes (Table 2 and S1 Fig). *Theileria parva* positive DNA samples from cattle in East Africa (Kenya, Uganda and Tanzania) and the non-clinical case from South Africa generated single amplicon profiles consisting of the 900 bp and 1000 bp fragments respectively (Table 2 and S1 Fig).

**Table 2.** PCR amplified p67 fragments detected in *T. parva* positive samples from cattle and buffalo

| Country |  | <sup>a</sup>Number of samples successfully amplified | <sup>b</sup>p67 fragment sizes detected in cattle |  |  |  | <sup>b</sup>p67 fragment sizes detected in buffalo |  |  |  |
| --- | --- | --- | --- | --- | --- | --- | --- | --- | --- | --- |
|  |  |  | 0.9Kb | 1.1Kb | 0.8Kb | 1Kb | 0.9Kb | 1.1Kb | 0.8Kb | 1Kb |
| Kenya | Nakuru Cattle | 12 | n=12 | ND | ND | ND |  |  |  |  |
|  | Olpejeta Buffalo | 22 |  |  |  |  | ND | n=22 | n=22 | ND |
| Uganda | Kiruhura cattle | 24 | n=24 | ND | ND | ND |  |  |  |  |
|  | Karamoja cattle | 06 | n=06 | ND | ND | ND |  |  |  |  |
| Tanzania | Tanga cattle | 03 | n=03 | ND | ND | ND |  |  |  |  |
|  | Simanjiro cattle | 06 | n=06 | ND | ND | ND |  |  |  |  |
|  | TNP Buffalo | 09 |  |  |  |  | ND | n=08 | n=09 | n=04 |
| South Africa | HIP Buffalo | 23 |  |  |  |  | n=04 | n=23 | n=23 | ND |
|  | CD clinical cases | 10 | n=04 | n=06 | n=06 | n=03 |  |  |  |  |
|  | Non-clinical *T. parva*-positive | 01 | ND | ND | ND | n=01 |  |  |  |  |
| Mozambique | MGR Buffalo | 12 |  |  |  |  | n=11 | n=8 | n=12 | n=3 |
| **Total** |  | **128** | **n=55** | **n=6** | **n=6** | **n=4** | **n=15** | **n=61** | **n=66** | **n=7** |

<sup>a</sup> TNP - Tarangire National Park; HIP - Hluhluwe-iMfolozi Park; MGR - Marromeu Game Reserve; CD - Corridor disease.
<sup>b</sup> ND - "Not Detected" for the respective allele.

### Detection of p67 allele types

A total of 230 p67 sequences were obtained, representing 85 samples selected for cloning (Table 3). The four p67 allele types were confirmed and the sequence identity was similar to the previously published and unpublished p67 sequences (Fig 1 and S6 Table). Allele type 1 sequences, which lack the predicted 43-amino-acid insert were common in all *T. parva* parasites from cattle from Kenya, Uganda and Tanzania, which comprised of both co-grazers (cattle that graze with buffalos) and non co-grazers, and the two vaccine stocks (Katete and Chitongo) from Zambia (Table 3 and Fig 1). In contrast, analysis of p67 sequences obtained from *T. parva* parasites from clinical cases of Corridor disease revealed representation of all four allele types (Table 3 and Fig 1). *Theileria parva* parasites from the single sample from non-clinical *T. parva*-positive case had only a single type sequence (allele type 4) (Table 3 and Fig 1). Notably, allele type 1 sequences were also identified from sequences from buffalo samples from southern Africa, but not in buffalo samples from East Africa (Table 3).

**Table 3.**
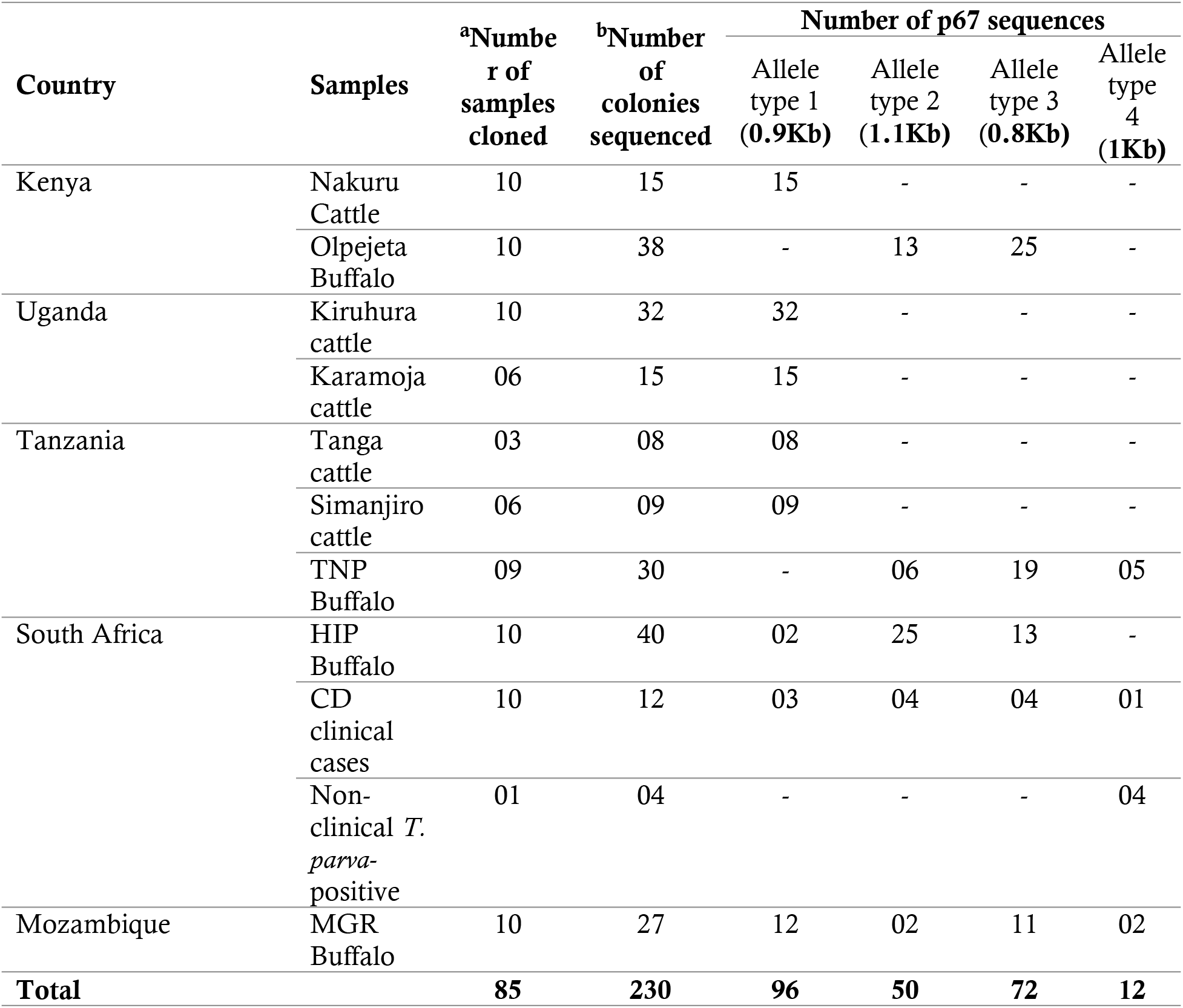
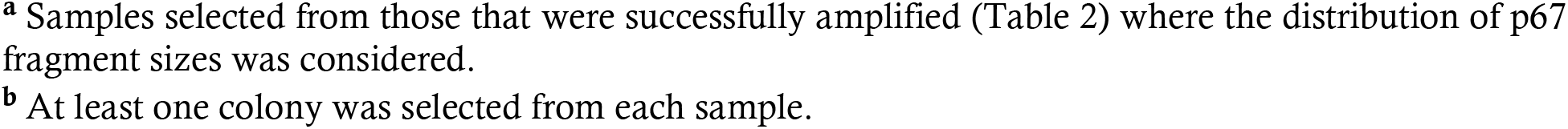
p67 allele types from *T. parva* positive samples from East and southern Africa

| Country | Samples | <sup>a</sup>Number of samples cloned | <sup>b</sup>Number of colonies sequenced | Number of p67 sequences |  |  |  |
| --- | --- | --- | --- | --- | --- | --- | --- |
|  |  |  |  | Allele type 1 (0.9Kb) | Allele type 2 (1.1Kb) | Allele type 3 (0.8Kb) | Allele type 4 (1Kb) |
| Kenya | Nakuru Cattle | 10 | 15 | 15 | - | - | - |
|  | Olpejeta Buffalo | 10 | 38 | - | 13 | 25 | - |
| Uganda | Kiruhura cattle | 10 | 32 | 32 | - | - | - |
|  | Karamoja cattle | 06 | 15 | 15 | - | - | - |
| Tanzania | Tanga cattle | 03 | 08 | 08 | - | - | - |
|  | Simanjiro cattle | 06 | 09 | 09 | - | - | - |
|  | TNP Buffalo | 09 | 30 | - | 06 | 19 | 05 |
| South Africa | HIP Buffalo | 10 | 40 | 02 | 25 | 13 | - |
|  | CD clinical cases | 10 | 12 | 03 | 04 | 04 | 01 |
|  | Non-clinical *T. parva*-positive | 01 | 04 | - | - | - | 04 |
| Mozambique | MGR Buffalo | 10 | 27 | 12 | 02 | 11 | 02 |
| **Total** |  | **85** | **230** | **96** | **50** | **72** | **12** |

<sup>a</sup> Samples selected from those that were successfully amplified (Table 2) where the distribution of p67 fragment sizes was considered.
<sup>b</sup> At least one colony was selected from each sample.

**Fig 1.**
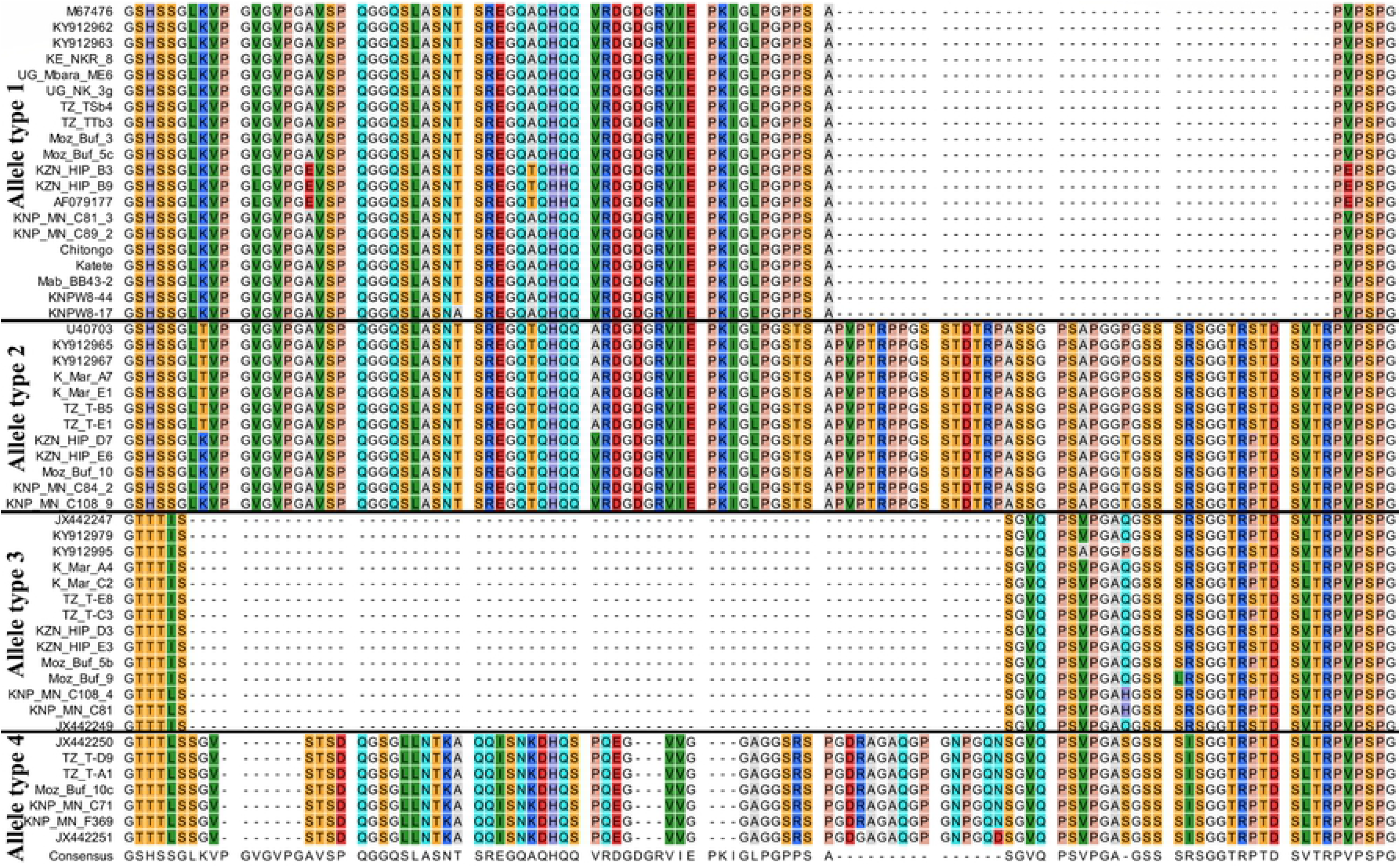
An alignment of p67 sequences from cattle- and buffalo-derived *T. parva* parasites. Kenya cattle (KE_NKR), Kenya buffalo (K_Mar), Uganda-Mbarara cattle (UG_Mbara), Uganda-Karamoja cattle (UG_NK), Tanzania-Tanga cattle (TZ_TT), Tanzania-Simanjiro cattle (TZ_TS), Tanzania buffalo (TZ_T), Mozambique buffalo (Moz_buf), KZN buffalo (KZN_HIP), CD clinical cases (KNP_MN_C), non-clinical *T. parva*-positive case (KNP_MN_F369) and reference sequences.

### Sequence variations in allele type 1

The p67 protein has two B-cell epitopes (TpM12 and AR22.7) recognized by murine monoclonal antibodies within the central variable region (19, 33). Analysis of the predicted protein sequences of the two epitopes in allele types 1, 2, 3 and 4 revealed amino acid substitutions in the buffalo-derived *T. parva* parasites, including those from clinical cases and the non-clinical *T. parva*-positive case from South Africa (Table 4 and S2-S4 Tables). Fourteen amino acid substitutions were detected in TpM12 epitope in all allele types from buffalo-derived parasites, while only one substitution was detected in AR22.7 epitope in allele type 2 (Table 4 and S2-S4 Tables). Out of the 14 substitutions in TpM12 epitope, allele type 1 had two substitutions at positions 170 and 183 (Table 4) in buffalo-derived *T. parva* parasites. However, both epitopes were conserved in allele type 1 detected from the cattle-derived *T. parva* parasites from Kenya, Uganda and Tanzania, and from the two isolates (Katete and Chitongo) from Zambia (Table 4).

**Table 4.**
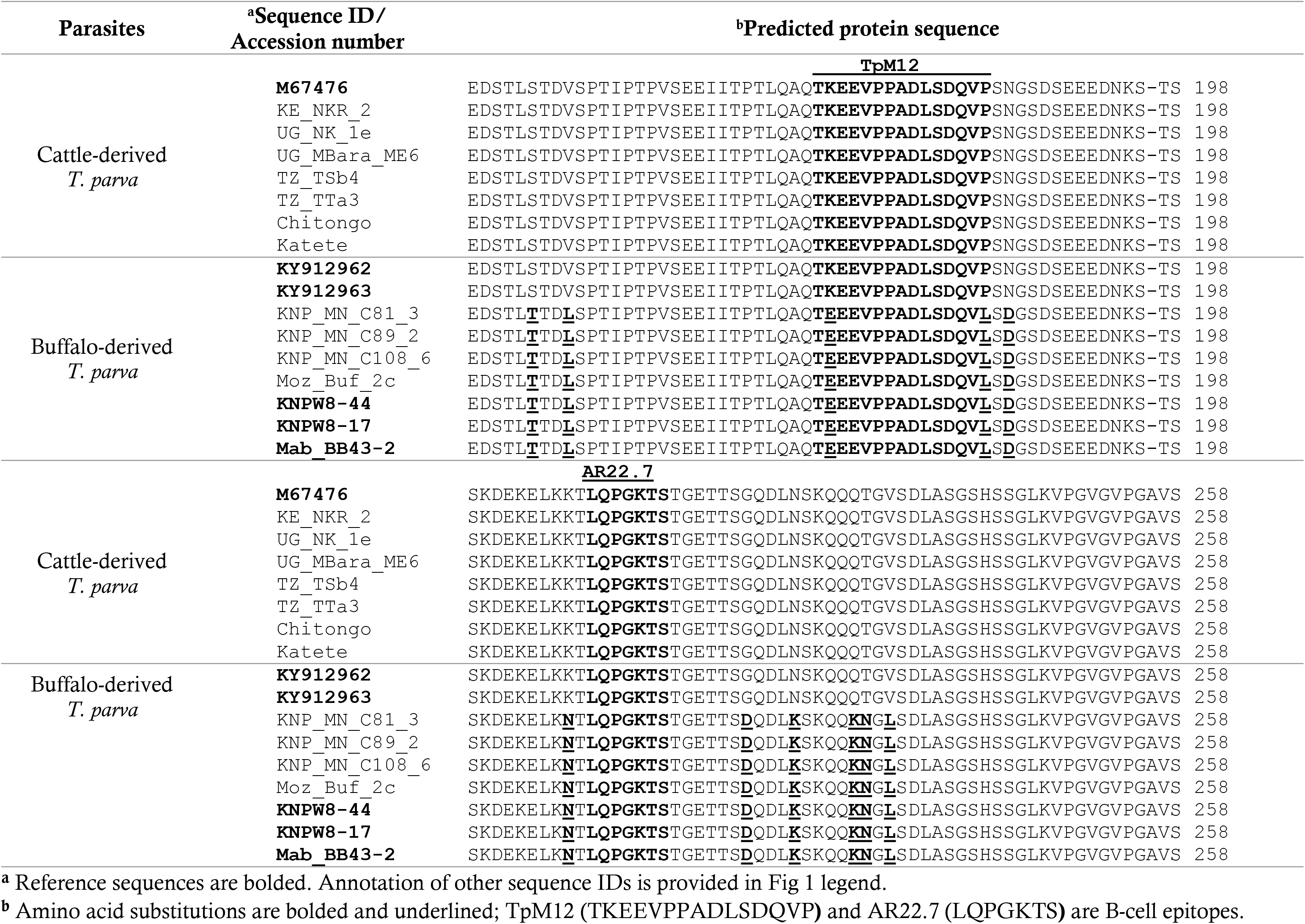
Predicted protein sequence alignment of allele type 1 showing sequence variations

Although the p67 allele type 1 was not detected from sequences from buffalo samples from East Africa in the current study, this allele type has been identified in parasites from buffalo in this region (34). Thus, allele type 1 predicted protein sequences obtained from cattle- and buffalo-derived *T. parva* parasites from East and southern Africa were compared, revealing two subtypes (Table 4). Allele type 1 subtype 1 sequences with 100% sequence identity to the Muguga isolate sequence were identified in cattle-derived *T. parva* parasites from Kenya, Uganda and Tanzania, and buffalo-derived *T. parva* parasites from cattle (KY912962) and buffalo (KY912963) in Kenya (Table 4). Allele type 1 subtype 2 sequences with amino acid substitutions at positions 170 and 183 in TpM12 epitope, and other unique substitutions within the amplified region were identified in buffalo-derived *T. parva* parasites from clinical cases of Corridor disease and buffalo from southern Africa (Table 4). The probability score for the prediction of the effect on the protein function as a result of the substitutions at positions 170 and 183 was 1.00.

### Phylogenetic analysis

Phylogenetic analysis of 34 randomly selected sequences representing the four p67 allele types and 18 reference sequences (ntax=52), comprised of 999 characters with 267 informative parsimony sites. GenBank accession numbers of some of the selected sequences from the current study are provided in S6 Table. Estimated proportion of invariant sites (p-inv) was 0.405 with Bayesian effective sample size (ESS) value >200 and a PhiTest (p-value) <1 (p=0.0) indicating significant evidence of recombination. Two statistically supported clades were recovered in all three analyses (Fig 2 and Fig 3). Clade A comprised of p67 sequences representing allele types 3 and 4, which were obtained from buffalo-derived *T. parva* parasites from buffalo from East and southern Africa, and cattle (Corridor disease cases) from South Africa (Fig 2). Clade B comprised of p67 sequences representing allele types 1 and 2 obtained from cattle- and buffalo-derived *T. parva* parasites (Fig 2). Among the allele type 1 sequences, there was a clear isolation pattern between *T. parva* parasites from East Africa (group 1) and southern Africa (group 2) (Fig 2). Allele type 1 group 1 comprised of sequences from cattle-derived *T. parva* parasites exclusively from East Africa, as well as sequences from buffalo-derived *T. parva* parasites from cattle (KY912962) and buffalo (KY912963) from Kenya. Allele type 1 group 2 comprised of sequences from the buffalo-derived *T. parva* parasites exclusively from cattle from South Africa, and buffalo from Kruger National Park in South Africa and Marromeu Game Reserve in Mozambique. Interestingly, from the network, allele type 1 group 1 and allele type 2 group 1, both from East Africa showed a strong genetic relationship and formed a cluster (Figure 3). Allele type 1 group 3 sequences from buffalo-derived parasites from Hluhluwe-iMfolozi Park in KwaZulu-Natal formed a small group that was slightly genetically dissimilar to the other allele type 1 groups, but with a close relation to allele type 1 group 2 sequences from parasites that circulate in and around Kruger National Park (Figure 3 and S5 Table).

**Fig 2.**
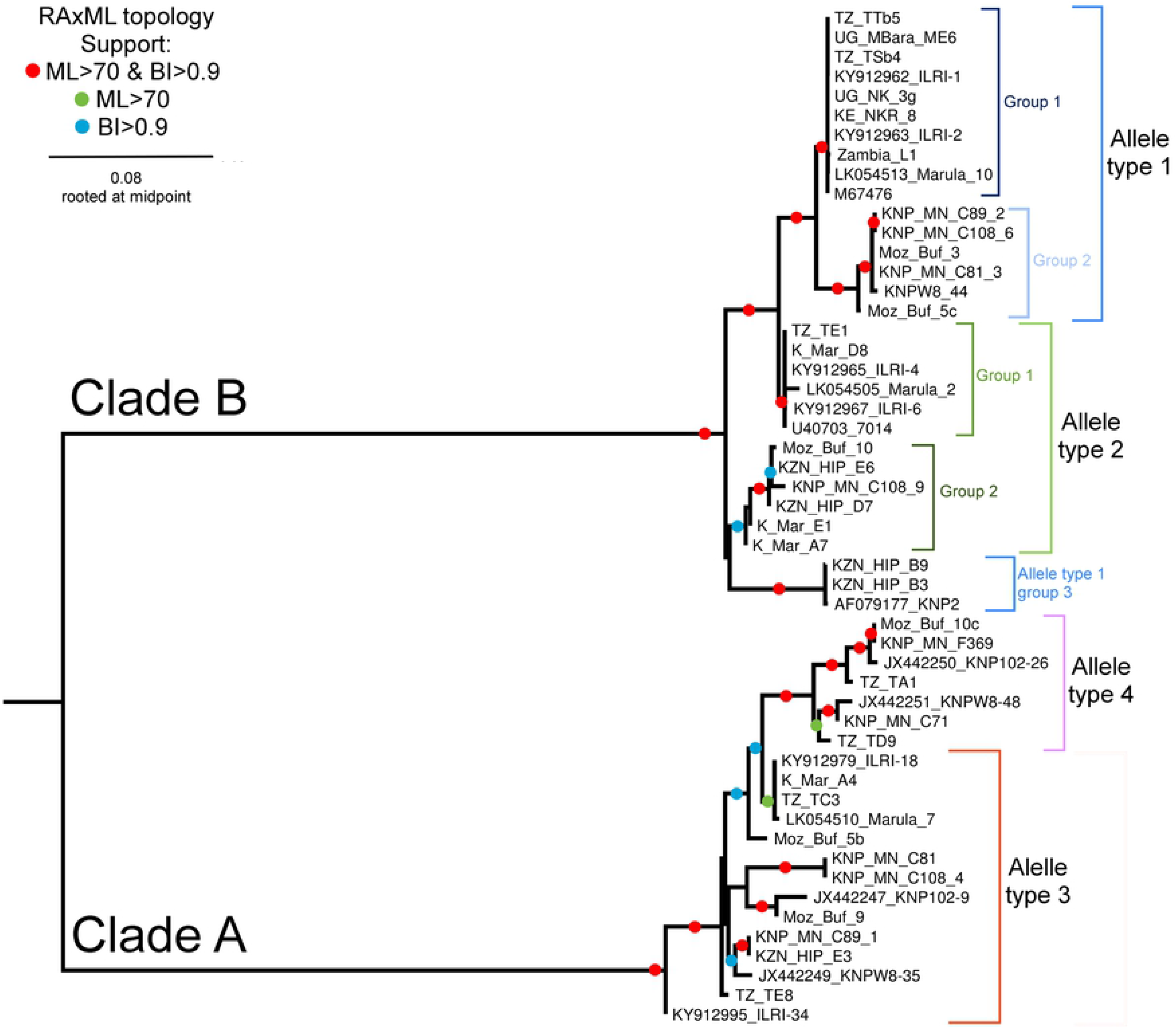
Topology recovered from maximum likelihood analysis in RAxML using GTR+I+G model of evolution. Support indicated on branches are bootstrap support from RAxML with autoMRE function invoked followed by posterior probabilities calculated from saved trees using Bayesian inference. Tree rooted at midpoint. Allele types are based on previous description (21, 26).

**Fig 3.**
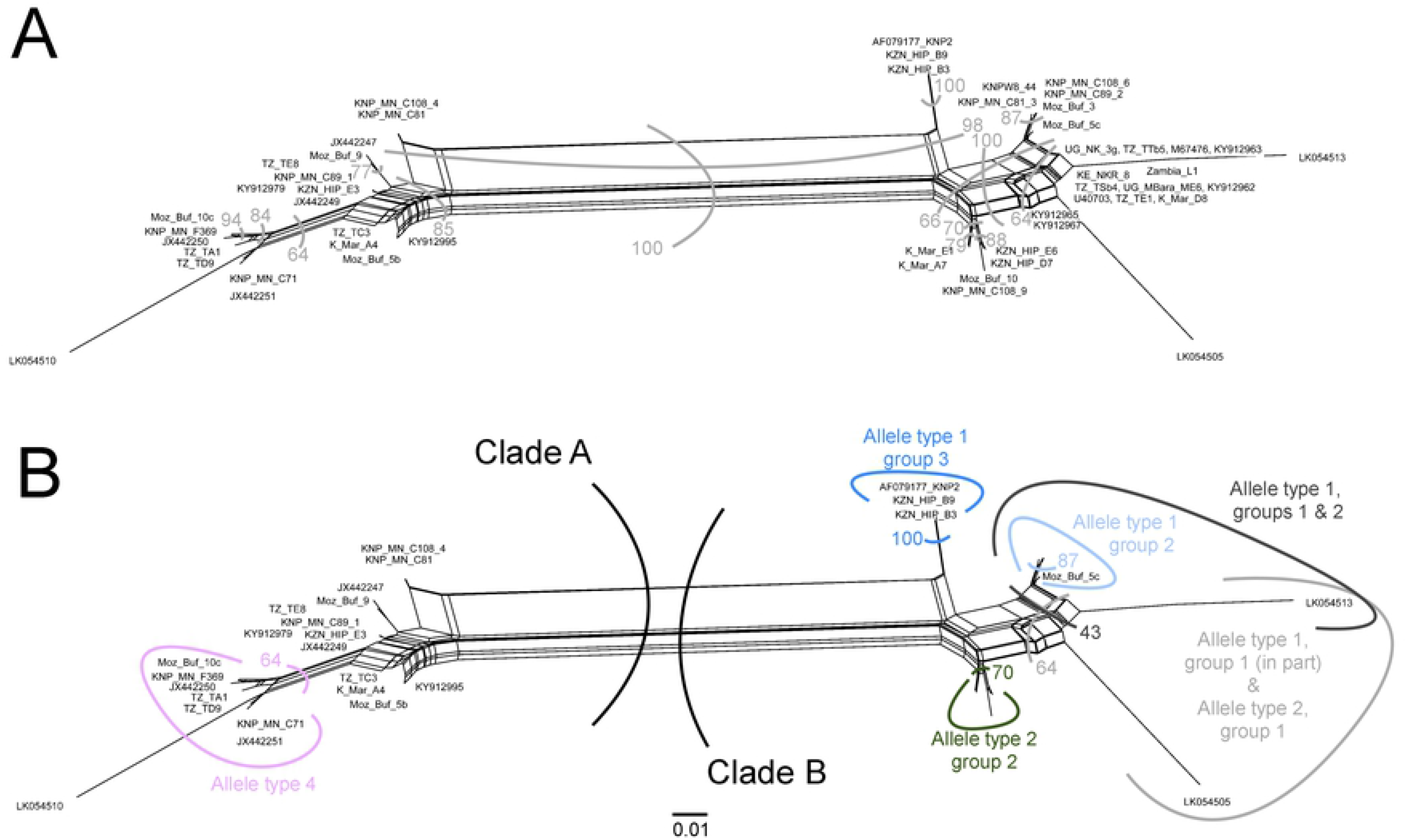
Data-display networks recovered from SplitsTree using all characters and uncorrected p-distances. Bootstrap support calculated from 1000 replicates. **A**: Indicates all major grouping with bootstrap support greater than 50. **B**: Allele types superimposed on network with accompanying support.

## Discussion

The central variable region of the p67 gene has been explored for assessment of the diversity of *T. parva* parasites originating from the buffalo and cattle in East and South Africa (19, 26, 33, 34). Due to the similarities in *T. parva* parasites on p67 allele type 1 and the associated disease syndromes in cattle in East and southern Africa, we evaluated cattle- and buffalo-derived *T. parva* parasites from the two regions to identify possible differences based on allele type 1.

Analysis of p67 amplicons and sequences from *T. parva* parasites obtained from cattle and buffalo blood samples from East and southern Africa revealed four allele types previously reported (21, 26). Buffalo-derived *T. parva* parasites from cattle in South Africa were heterologous with all the four p67 allele types present while cattle-derived *T. parva* parasites from East Africa (Kenya, Uganda and Tanzania) were invariant with only allele type 1 identified. Collectively, the results of the current study and previous reports (21, 33) demonstrate a remarkable reduction in the diversity of the p67 gene in cattle-derived *T. parva* parasites in the study areas in East Africa. A conceivable explanation to this could be that a subpopulation of *T. parva* parasites from the buffalo probably possessing allele type 1 have undergone selection and adaptation for cattle to cattle transmission, and are circulating in the cattle population in eastern Africa.

We established that the p67 allele type 1 sequences from *T. parva* parasites transmitted naturally from the buffalo to cattle in Kenya (34), as well as from parasites from cattle that co-graze with buffalo in Mbarara-Uganda and Simanjiro-Tanzania were identical to those from the cattle-derived parasites from East Africa. *Theileria parva* parasites possessing p67 allele type 1 have been associated with classical ECF in eastern Africa (21). From the phylogenetic analysis, it is likely that parasites possessing allele type 1 evolved from the buffalo-derived parasites possessing allele type 2, and have adapted for cattle-to-cattle transmission (45). The former could be the subpopulation of parasites that is circulating in cattle in East Africa, which can establish a carrier state more efficiently in cattle in the region, and responsible for classical ECF. To further establish any possible genetic alterations associated with the presumed adaptation event, it could be of interest to sequence and compare the genomes of p67 type 1 and 2 parasites derived from the buffalo, and type 1 derived from *T. parva* parasites circulating in cattle in East Africa.

Sequence analysis of two p67 epitopes (TpM12 and AR22.7) reported to be reactive to murine monoclonal antibodies (19), and are also a target of the host’s B-cell responses (33), revealed a subtype of p67 allele type 1 from the buffalo-derived *T. parva* parasites from clinical cases of Corridor disease in South Africa. This finding indicates that not all parasites possessing allele type 1 are associated with classical ECF, which has not been reported in South Africa since its eradication in the early 1950s. *Theileria parva* parasites possessing this subtype could be part of the parasites that cannot establish a long-lasting carrier state in cattle, and possibly responsible for Corridor disease in South Africa. It is worth noting that the role of parasites possessing allele type 1 in the pathogenesis of Corridor disease in East and southern Africa still remains unclear. The clinical presentation of Corridor disease in South Africa (46) and East Africa (13) is similar, yet some of the parasites involved differ based on p67 allele type 1. Therefore, it is most likely that the differences between the parasites involved, based on p67 allele type 1, have no link to disease manifestation, but rather suggestive of confinement of the parasites possessing the respective allele type 1 subtypes to the two regions. Nevertheless, it was recently established that the diversity of *T. parva* parasites arose before geographic separation of eastern and South African parasites (47), and the predominance of p67 allele types in parasites from the two regions vary (26, 33, 34). Thus, it is possible that both subtypes of p67 allele type 1 are present in the two regions, but some are underrepresented in one geographical region over the other.

Corridor disease samples from South Africa were obtained from the designated Corridor disease infected areas bordering Kruger National Park in Mpumalanga province and Hluhluwe-iMfolozi Park in KwaZulu-Natal province. Due to very low parasitemia, we were not able to generate p67 amplicons from the four cattle samples from uMkhanyakude and Hluhluwe districts that were positive for *T. parva* on real-time PCR. Therefore, to further support the results of the current study, it would be necessary to analyze more samples from Corridor disease cases in KwaZulu-Natal where the disease is likely to occur. Interestingly, two allele type 1 sequences obtained from parasites from buffalo from Hluhluwe-iMfolozi Park were slightly different (with ~ 6% sequence difference) from the other type 1 sequences from South Africa. This indicates that although allele type 1 sequences have a defining 129 bp indel associated with them, there could be more polymorphism in this allele type in parasites from southern Africa. Nonetheless, the results of the current study form the first report on the difference in the genotypes of *T. parva* parasites from East and southern Africa, based on p67 type 1 allelic sequences.

A subtype of p67 allele type 1 identical to that identified from parasites from clinical cases of Corridor disease from South Africa was also detected from *T. parva* parasites from the buffalo from Marromeu Game Reserve (MGR) in Mozambique. The other allele types (2, 3 and 4) were also similar between the two populations from southern Africa. The buffalo herd in MGR has been isolated for many years in the Zambezi delta within the East coast of southern Africa, and there is no official report of Corridor disease in the area adjoining MGR. Thus, assuming that the subtype of p67 allele type 1 identified plays a role in the resulting disease, the results suggest that should there be contact between the Zambezi delta buffalo herd and naïve cattle in the presence of the tick vector, there is a likelihood that an outbreak of Corridor disease could occur.

The phylogenetic trees and network generated, displayed congruent clustering comparable to analyses conducted previously (26). Notably, the clustering separated the subtype of allele type 1 from parasites associated with Corridor disease in South Africa, from the subtype detected from parasites associated with ECF in East Africa and also detected in parasites originating from buffalo in Kenya. These findings on the evolution of *T. parva* provide additional evidence on the difference between parasites from the two regions of Africa. Therefore, the concern about the possibility of re-emergence of ECF in South Africa based on p67 allele type 1 (26) may be annulled. Nonetheless, should the buffalo-derived *T. parva* adapt to cattle to establish a long-lasting carrier state, then this scenario poses a high risk to the cattle population in South Africa. Currently, the movement of buffalo is strictly monitored in South Africa by the South African Directorate of Animal Health (Veterinary Procedural Notice: Buffalo Disease Risk Management), in addition to the general cattle-buffalo isolation practices, hence reducing the possibility of transmission of parasites, including *T. parva*, between the two species.

We compared the variation of *T. parva* parasites from East and southern Africa on the two p67 epitopes. The predicted protein sequences of the subtype of p67 allele type 1 identified from the buffalo-derived *T. parva* parasites from southern Africa had two amino acid substitutions at positions 2 and 15 in TpM12 epitope (T<u>**E**</u>EEVPPADLSDQV<u>**L**</u>) and none in AR22.7. The probability that these amino acid substitutions in TpM12 would significantly alter the structure and antigenicity of the epitope is low (probability score = 1.00). However, further investigation will be necessary to ascertain whether these variations affect recognition by murine monoclonal antibodies. The effectiveness of the current p67 recombinant vaccine in the field is dependent on its ability to induce production of sporozoite neutralizing antibodies that are protective against a homologous challenge with *T. parva* sporozoites (19). Notwithstanding that, the vaccine confers partial protection in the field against a challenge with ticks infected with cattle-derived *T. parva* (31). Although it has been suggested that inclusion of additional sporozoite surface antigens could improve the p67-based recombinant vaccine (48), protection against a sporozoite challenge with buffalo-derived *T. parva* parasites may continue to be a challenge, especially with the B-cell epitope variations reported in buffalo-derived *T. parva* parasites from cattle in the current study.

## Conclusion

We report identification of a subtype of p67 allele type 1, unique to buffalo-derived *T. parva* parasites from southern Africa, which can be explored as a marker to differentiate *T. parva* parasites of p67 allele type 1 responsible for ECF and Corridor disease. Further analysis of more samples from Corridor disease cases and buffalo from southern Africa is recommended to support this finding.

## Supporting Information

**S1 Fig.** PCR amplicons from cattle- and buffalo-derived *T. parva* parasites from East and southern Africa.

**(a)** p67 PCR amplicons from buffalo-derived *T. parva* parasites from clinical cases of Corridor disease (CD) and non-clinical *T. parva*-positive cases from South Africa (SA); **(b)** p67 PCR amplicons from buffalo-derived *T. parva* parasites originating from buffalo in Hluhluwe-iMfolozi Park, KwaZulu-Natal; **(c)** p67 PCR amplicons from cattle-derived *T. parva* parasites originating from cattle in Mbarara district in Western Uganda. 1kb DNA ladder (#SM0311, ThermoFisher Scientific, Waltham, MA USA) was used in (a), 100bp plus DNA ladder (#SM0321, ThermoFisher Scientific, Waltham, MA USA) was used in (b) and (c). SA – South Africa; CD – Corridor disease; M – molecular weight marker.

**S1 Table.** The Ct values of samples from active clinical cases of Corridor disease and non-clinical *T. parva*-positive cases collected from Mpumalanga province in South Africa.

**S2 Table.** Predicted protein sequence alignment of allele type 2 identified in *T. parva* parasites from cattle and buffalo.

**S3 Table.** Predicted protein sequence alignment of allele type 3 identified in *T. parva* parasites from cattle and buffalo.

**S4 Table.** Predicted protein sequence alignment of allele type 4 identified in *T. parva* parasites from cattle and buffalo.

**S5 Table.** Estimates of the evolutionary divergence between sequences of allele type 1 from *T. parva* parasites from East and Southern Africa.

**S6 Table.** Taxonomic metadata detailing the grouping of p67 allele types from *T. parva* parasites from East and Southern Africa.

## Acknowledgments

The authors wish to thank SANParks (South Africa) for providing buffalo samples, Nicola Collins and Luis Neves for providing cattle DNA samples from South Africa and buffalo DNA samples from Mozambique respectively, Isaiah Obara for providing buffalo DNA samples from Kenya, and Paul Gwakisa for providing buffalo and cattle DNA samples from Tanzania.

